# Can seminal fluid proteins be honest signals of sperm availability?

**DOI:** 10.1101/2024.04.26.591243

**Authors:** Piotr Michalak, David Duneau, Jean-Baptiste Ferdy

**Affiliations:** Centre de Recherche sur la Biodiversité et l’Environnement - CRBE UMR5300, Université de Toulouse, CNRS, IRD, Toulouse INP, Université Toulouse 3 – Paul Sabatier (UT3), Toulouse, France; The Queen’s Medical Research Institute, Centre for Cardiovascular Science, Edinburgh University, Edinburgh, UK

**Author notes:** Corresponding author: Piotr Michalak. these authors have contributed equally (David Duneau, Jean-Baptiste Ferdy).

**Keywords:** signals, oviposition, seminal fluid, proteins, drosophila, sexual conflict

## Abstract

Seminal fluid proteins (Sfps) are produced by males, yet they have significant effects on female physiology and behaviour. Sfps are often viewed as a mechanism for males to manipulate female reproductive behaviours, but despite many studies identifying their varied effects and their molecular underpinnings, the ways males and females use Sfps remain unclear. In the most studied case of *Drosophila melanogaster* particular peptides within the seminal fluid have been identified to guide female reproduction: sex peptide (SP, Acp70A) is known to induce changes in egg production, oviposition and remating and is attached to sperm and continuously supplied for days after mating, while ovulation inducing peptide, ovulin (Acp26Aa), is a free peptide with only short term effects after mating. Using a biologically informed model we show how these peptides can synchronize sperm and egg release, hence reducing the number of unfertilized eggs a female lays. We further show that the exhaustion of SP might be the key signal of this synchronization. Finally, we demonstrate that sexual conflict over the regulation of female reproductive physiology by Sfps should be limited, with the primary conflict probably centring around the regulation of remating behaviour.

## Introduction

The signals exchanged between males and females before and during mating have been extensively studied in the context of sexual selection (Andersson, 1994; Maynard Smith & Harper, 2003; Dougherty, 2021; Rosenthal & Ryan, 2022). Among the different signals exchanged, seminal fluid proteins (Sfps) are unique: they are provided by the male, persist but change after mating (Peng et al., 2005) and can influence female behaviour long after the male has left. Sfps are found across different animal groups, including nematodes, insects, crustaceans, birds, mice and humans, and have a range of effects on female reproductive behaviour, physiology and immunity (Ward & Carrel, 1979; Fujihara, 1992; Eberhard, 1996; Schjenken & Robertson, 2020; Guan et al., 2023). Despite widespread occurrence and functional similarity of Sfps, they often lack common ancestry and show high rates of evolution (Sirot et al., 2015; Hopkins & Perry, 2022; Hurtado et al., 2022). This fast divergence, their male origin, the profound changes they induce in the mated females physiology and behaviour as well as their induced fitness costs have led to the view of Sfps as potent weapons in the sexual conflict (Sirot et al., 2015).

Seminal fluid proteins have been extensively studied in insects, especially in *Drosophila melanogaster*. In *Drosophila*, the changes after mating have been found to be driven by sex peptide (SP, Acp70A), a peptide produced in male accessory gland (Liu & Kubli, 2003). SP is provided along the ejaculate and is bound to the sperm tail in the female reproductive tract (Chen et al., 1988; Peng et al., 2005; Misra et al., 2022). After mating, when sperm is stored, SP is cleaved away from sperm and released into female reproductive tract, binding to SP receptors and passing a signal to the brain (Peng et al., 2005; Yapici et al., 2008; Yang et al., 2009; Rezával et al., 2012; Kubli & Bopp, 2012; Yang et al., 2023). The consequence is an increase in female activity, aggression, a reduction of the rest periods, a reduction in remating (Hopkins & Perry, 2022; Yang et al., 2023). SP also increases the synthesis of the juvenile hormone (JH) and stimulates egg production and oocyte maturation (Moshitzky et al., 1996; Berg et al., 2024). Finally, SP increases the oviposition and the use of sperm from the female storage compartments (Chapman, 2001; Ravi Ram & Wolfner, 2007; Avila et al., 2015; Hopkins & Perry, 2022). These long-term effects are sustained by the continuous release of sperm bound SP for several days (Peng et al., 2005). Although SP increases egg production, it can also impose costs on females: SP lowers resistance to infection, decreases survival, and can even lower fecundity of females continuously exposed to it (Schwenke et al., 2016; Schwenke & Lazzaro, 2017; Chapman et al., 1995; Wigby & Chapman, 2005). Because of these costs, SP has become a staple example of sexual conflict and of male manipulation (Chapman et al., 1995; Arnqvist & Rowe, 2005; Wigby & Chapman, 2005). However, a recent review suggests that a purely antagonistic role of SP is unlikely and that neutral or positive effects for the female are more supported by the evidence (Hopkins & Perry, 2022).

The post-mating response is usually regulated by several different Sfps, often overlapping in function (Eberhard, 1996). In *Drosophila*, ovulin, (Acp26Aa), is provided in the form of free peptides in the seminal fluid. After mating, ovulin is found primarily at the base of the ovary and increases ovulation and oviposition by the relaxation on the reproductive tract musculature (Monsma & Wolfner, 1988; Herndon & Wolfner, 1995; Heifetz et al., 2000; Rubinstein & Wolfner, 2013). The ovulin induced increase in oviposition is short term, ending within 24 hours after mating (Herndon & Wolfner, 1995). This might benefit both or only one of the sexes, but there are no known significant effects of ovulin on female fitness. The potential of ovulin in sexual conflict has been only briefly sketched and remains largely unexplored (Herndon & Wolfner, 1995; Heifetz et al., 2000; Sirot et al., 2015).

Sfps influence many aspects of female reproduction, making it difficult to determine which effect prevails. SP and ovulin have been proposed as means of aligning male-female interests, but also as a way of male manipulation (Hopkins & Perry, 2022; Sirot et al., 2015). Dissecting the different functions of SP and ovulin to understand their fitness consequences is intricate, and we lack clear predictions to guide experimental approaches. We developed a mathematical model of female reproductive physiology and of its regulation by Sfps. Our model is based on the reproductive physiology of *Drosophila melanogaster*, but the logic applies broadly to sperm storing insects. We focused on how the SP and ovulin influence egg production, as well as oviposition and sperm usage and studied the trade-offs between three fitness relevant proxies obtained from the model. We show how female might use signals from Sfps to guide her reproductive decisions and reduce fitness costs. We also show the conditions under which male and female interests diverge and that potential for conflict raises largely when females remate.

## Fitness trade-offs under unregulated reproduction

### The model without regulation of egg laying

To determine the role of Sfps in female reproduction, we started by studying what happens in the absence of any signal. We assumed that sperm transfer does not change female behaviour; she lays a constant, maximal number of eggs. Gilbert et al. (1981) proposed a statistical, discrete time model of *Drosophila melanogaster* reproduction without regulation. They assumed that the number of fertilized eggs laid is driven by whichever gamete type, sperm or eggs, is limiting. We have made the same assumption, but supposed that egg production and fertilization can happen at any time. We thus considered time as being continuous and used differential equations (Fig. 1).

**Figure 1.**
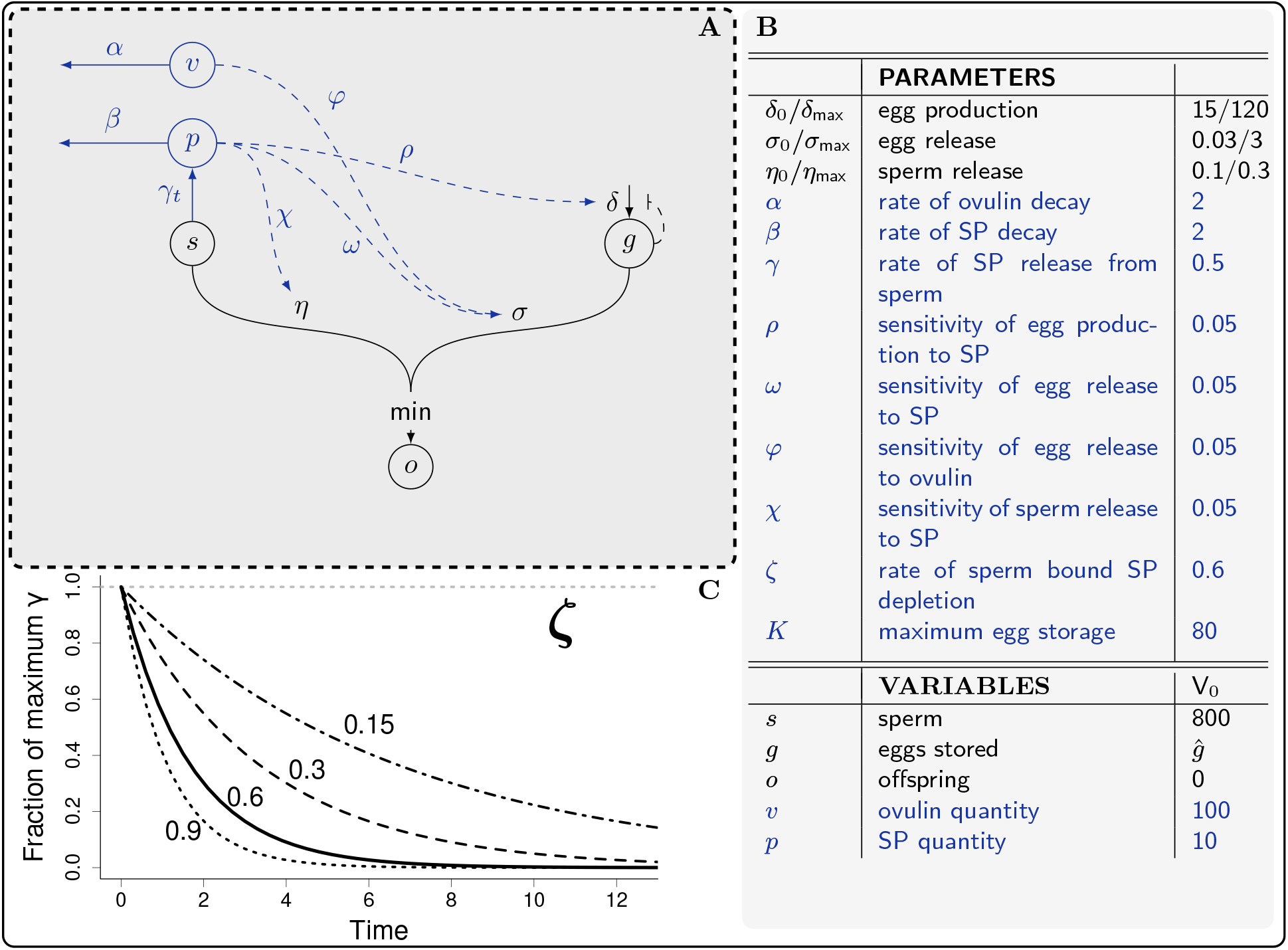
(A) Schematic of the model of reproduction. Without regulation by Sfps (black), available sperm (s) and eggs (g) follow independent dynamics. Under regulation by Sfps (blue), numbers of released sperm and eggs that determine offspring production are connected through SP (*p*) supplied from sperm. (B) Parameters’ values and initial state of the variables used in our simulations, except when otherwise stated. 0 and max subscripts correspond to base and max values, respectively. (C) SP on sperm decreases over time. Unless there is substantial delay in SP decay, SP detachment rate determines the end of SP activity.

Adult females produce eggs that are stored up to a genetically and developmentally determined capacity, which in our model, we called *K*. The eggs are produced at a rate related to the number of ovarioles (Flatt, 2020) and after a number of eggs have been stored, the production slows down and further oocytes are resorbed (Ashburner, 1989). We considered that the stored eggs are released at a constant rate, so the change in how many eggs are stored over time is described by

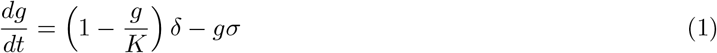

where *g* is the number of stored eggs, *δ* the maximum number of eggs produced per unit of time and *σ* the rate at which eggs are released. Over time, the number of eggs in storage approaches the equilibrium capacity determined by the balance between egg production and egg release:

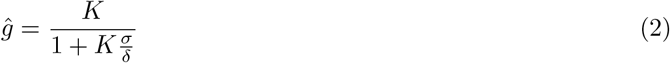

which equals the maximum capacity ĝ = *K* if egg release is blocked (*σ* = 0) and decreases when the ratio between egg release and egg production (*σ*/*δ*) increases.

To determine specific parameters’ values we follow the biology of *Drosophila*. The maximum number of stored eggs (*K*) is difficult to quantify, since females lay eggs even when not mated. On average, females can retain two eggs per ovariole in their post-reproductive period, when they cease oviposition (King, 1970; Klepsatel et al., 2013). Female fly has approximately 30 to 40 ovarioles (Boulétreau, 1978; Klepsatel et al., 2013), thus we considered a maximum capacity of *K* = 80. While we cannot exclude the possibility that young females may retain more eggs than older ones, this estimate is likely within a biologically plausible range.

We aimed to parameterize the model so that one unit of time corresponds to roughly one day in fly lifetime. *δ* is the maximum number of eggs that can be produced within a day. The actual production of eggs per day in the model is 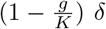 which depends on the current value of eggs in storage (*g*). The production of eggs is therefore connected to the rate of egg release: higher egg release lowers the number of eggs in storage, increasing egg production. This interdependence means that obtaining the values of egg production parameter (*δ*) is not simply equivalent to the experimental observations of oocyte maturation. We employed equation (2) to hypothesize about values approximating the real-world scenario. In the first ten days of reproduction, when reproductive output is maximal, flies laid daily approximately two eggs per ovariole (Klepsatel et al., 2013). We thus posit that the number of released eggs (*ĝσ*) is twice the number of ovarioles. Using our estimate of *K* = 80 for the assumed 40 ovarioles, we searched through values of egg production (*δ*), release (*σ*) and equilibrium egg storage (*ĝ*) that match these conditions. We propose *δ* = 120 and *σ* = 3, which yields *ĝ ≈* 26.6 eggs in storage at the equilibrium, a value within the experimentally observed 0-50 range (Boulétreau, 1978). At mating (*t* = 0) females store sperm and release it gradually to fertilize the eggs (Bloch Qazi & Wolfner, 2006). We describe this by:

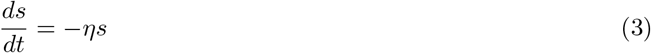

where *s* is the number of individual sperm cells stored and *η* is the rate of sperm release. The number of sperm a female store at any time can be obtained as the solution of (3):

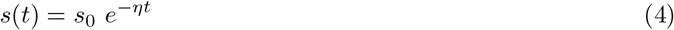

where *s*_0_ is the number of sperm stored at the time of mating, limited by the storage capacity. We used *s*_0_ = 800 based on the experimentally observed range (Bloch Qazi & Hogdal, 2010). In the same experiments, sperm depletion happened around 10 or 15 days post mating, depending on new egg laying substrate frequency, and 50 % of sperm was lost in 3-5 days. Sperm release parameter (*η*) in range from 0.1 to 0.66 would be consistent with those observations (we ignore the substrate dependence to keep the model simple and facilitate its interpretation). Unlike egg production or release, sperm release lacks a clear definition of maximal reproductive effort. As sperm number is limited in the model, very fast sperm depletion might lead to a rapid loss of sperm, without increasing the number of offspring. Unless otherwise specified, we thus have used an intermediate value of sperm release parameter of *η* = 0.3, which means that 95 % of sperm is used within ten days.

The number of produced offspring per unit time is assumed to be equivalent to whichever gamete released is limiting (Gilbert et al., 1981) which yields

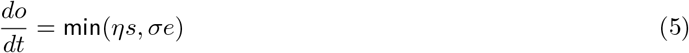

where *o* is the number of offspring produced since the time of mating. Because sperm and eggs equations (equations (1) and (4)) are independent and tractable, we can obtain a full solution for the total offspring production:

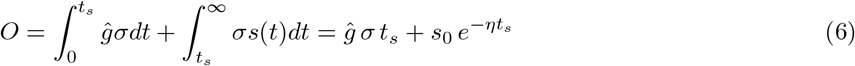

In our calculation the number of offspring is obtained as the number of released eggs until sperm becomes limiting, plus the number of sperm left in storage at that time. Equation (6) assumes that the female has reached a stable state of eggs in storage before mating (i.e. *g*(0) = *ĝ*) and that just after mating the production of offspring is limited by eggs, rather than by sperm. When this assumption holds, sperm becomes limiting at a time *t*_*s*_ defined by *σg*(*t*_*s*_) = *ηs*(*t*_*s*_). If we relax this assumption (thus *σĝ > ηs*_0_) the total offspring is maximal and simply equals the number of sperm provided (*O* = *s*_0_). Total offspring production should be understood as a theoretical value, corresponding to the lifetime offspring production a female fly would have if mating only once and without ageing. This value is an indication of the efficiency of reproductive physiology and avoids complications of multiple matings occurring at time intervals that depend on additional factors.

The dynamics which correspond to equations (1)-(6) are shown in Fig. 2A-B. As her physiology is not regulated, the female releases a constant, maximal number of eggs throughout. This results in a linear increase in the offspring number from mating onwards, as long as the number of released sperm exceeds that of released eggs (Fig. 2B). As sperm gradually depletes, the rate of sperm release falls below that of egg release at *t* = *t*_*s*_ (Fig. 2A). Offspring production then slows down, as its rate follows sperm release, until the number of produced offspring approaches a plateau (Fig. 2B). As the female consistently lays a maximum number of eggs per unit of time, the cumulative count of unfertilized, and therefore wasted, eggs that she lays continues to increase.

**Figure 2.**
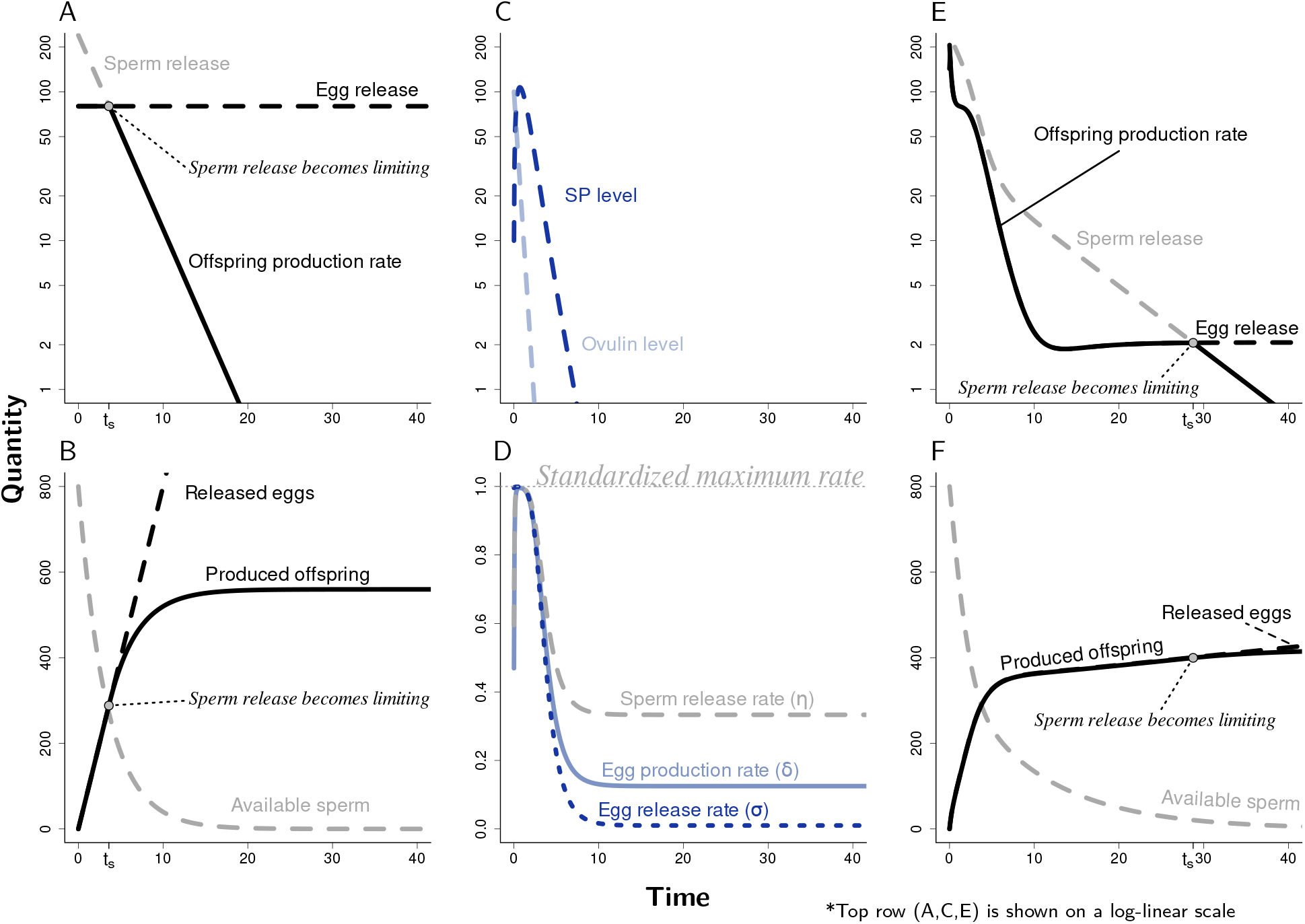
Simulation of female reproduction without regulation. (A) Instantaneous rates for three offspring determining processes - sperm release, egg release and offspring production. At time of mating (t=0), egg storage is at equilibrium. Egg release remains constant. Sperm release is initially high but slows down. The gamete type released in lower quantity drives offspring production. (B) Cumulative quantities from the curves in (A). As sperm becomes depleted, offspring production slows down and plateau near the final total offspring. The eggs are continuously produced at the same rate, even after the sperm becomes limiting (*t*_*s*_) leading to wastage of eggs. **Simulation of female reproduction with regulation.** (C) Ovulin declines exponentially over time. SP initially increases and eventually declines exponentially. (D) Effect of ovulin and SP on three main regulatory functions. Sperm release (*η*) and egg production (*δ*) rates are synchronous and exhibit an early peak matching SP’s, egg release rate(*σ*) does not. (E) Rates over time. Egg release exhibits an early peak driven by ovulin and increasing SP. When SP levels decline, egg and sperm release slow down, delaying sperm limitation (*t*_*s*_) to 10-fold. (F) Cumulative quantities from the curves in (E). Offspring production closely follows the pattern of egg release.

There is no single set of parameters that corresponds to real life scenario that we could identify. Therefore, we studied the impact of parameter variation, first focusing on sperm and egg releases (*η* and *σ*). Those two parameters are the main determinant of the number of produced offspring, since they are responsible for the number of available gametes (5). They are also not interconnected, unlike egg production and egg release (see (1) and (2)) and thus influence different processes of the reproductive strategy. To measure their influence, we computed three quantities which summarize the dynamics shown in Fig. 2A-B, and which can be considered as proxies of female fitness: total number of offspring, the time to reach 50 % of the total offspring, and the number of unfertilized eggs when 99 % of offspring has been reached. Total offspring production (*O* in (6)) is similar to *R*_0_, and its connection to fitness is immediate. The time a female takes to produce offspring is also part of her fitness: slow reproduction poses the risk of death before reaching maximal reproductive output. Finally, the number of unfertilized eggs provides a crude estimate of the energy efficiency of a particular reproductive strategy, as each unfertilized egg is a unit of energy that has been wasted and is not available for offspring production. The number of offspring only approaches its maximum as time goes to infinity, while the number of unfertilized eggs exhibits a linear and indefinite increase. Thus a meaningful number of unfertilized eggs can only be computed on a selected time frame and we opted to calculate it when females have reached 99 % of their total offspring production.

### A constant rate of oviposition leads to trade-offs between total offspring production and egg wastage

We now turn to comparison of different strategies of sperm and egg release and its fitness consequences. As mentioned before, the total number of offspring is at maximum when initial sperm release is below that of egg release (*s*_0_*η ≤ σĝ*). This happens left of the red curve on Fig. 3: all the strategies of egg and sperm release in this parameter range yield maximum offspring production, as number of released eggs can never fall below that of sperm. However, if the number of released sperm is higher than the number of released eggs at the time of mating (*s*_0_*η > σĝ*) some sperm is wasted and the total number of offspring is below the possible maximal value (*O < s*_0_) (which happens right of the red curve, Fig. 3A). Thus releasing sperm too fast tends to decrease total number of offspring, while it is maximized when sperm release is extremely slow. Here female can continuously produce and release eggs.

**Figure 3.**
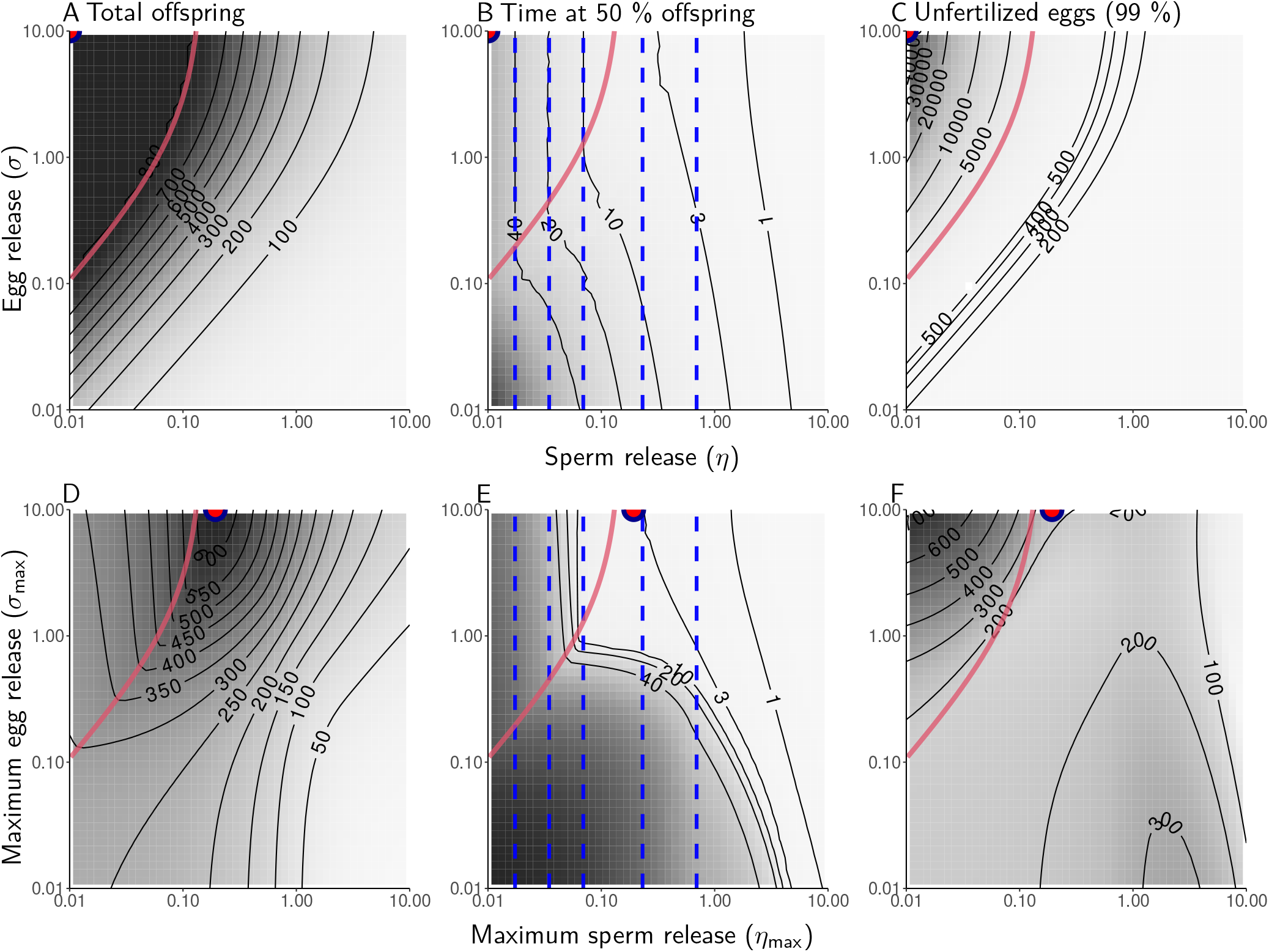
Panels present the three components of fitness for varying values of sperm and egg release parameters. In the model with regulation *η* and *σ* are not fixed and cannot be arbitrarily varied, so maximal values are used. **Model without regulation necessitates fitness trade-offs (A-C)** The red-blue point is an example of an offspring maximizing strategy. Total offspring production (A) cannot be maximized without reproducing slower (B, black contour lines show Time to reach 50 % of total offspring, blue dashed lines time to it takes to release 50 % of sperm in storage) and without wasting eggs (C). **Regulation by sex peptide relaxes trade-offs between fitness components (D-F)** The point maximizing offspring production (D) is in space of fast reproduction (E) and lower egg wastage (F).

If the sperm is always limiting, the time at which 50 % of offspring is produced (the black contour lines in Fig. 3B) depends entirely on the rate of sperm release and is simply that at which 50 % of stored sperm has been released, thus corresponding to the vertical dashed blue lines. This happens only left of the red curve (Fig. 3B) i.e. when released sperm is always limiting. To the right of the curve, when sperm is initially released in excess, black contour lines depart from blue dashed lines as the 50 % of offspring is reached later, likely because the early reproduction consist a smaller fraction of the total offspring production. In the model, sperm depletion puts an end to reproduction and faster sperm release results in faster reproductive dynamics.

Because the number of released eggs remains constant, there is always fewer sperm released late in the reproductive period, increasing the proportion of unfertilized eggs. This loss is largest when sperm is always limiting (left of the red curve, 3C), suggesting that the slow sperm release that maximizes total offspring production (Fig. 3A) also leads to the wastage of large quantities of eggs.

Overall, a strategy that would lead to the maximal number of offspring (such as the red-blue point in Fig. 3A-C), is a strategy where reproduction is very slow and where many eggs are wasted: in the absence of regulation, there is a trade-off between the maximal production of offspring and other fitness components. This happens because of a mismatch between the dynamics of egg and sperm, yielding periods where one largely exceeds the other so that either male or female gametes are wasted.

## Fitness trade-offs under regulation by seminal fluids

### The model with regulation by seminal fluid proteins

We found that the mismatch between the numbers of released eggs and sperm creates trade-offs between the different components of fitness. We hypothesized that the regulation of reproduction by Sfps could coordinate egg and sperm release and relax these trade-offs. Among *Drosophila* Sfps the peptide primarily responsible for regulation of modelled processes (egg production, release and the release of sperm from storage) is sex peptide (SP). Another peptide, ovulin, is responsible solely for the increased ovulation. We consider that SP (*p*) is provided both in the form of free peptides in the seminal fluid, and in the sperm bound form (Peng et al., 2005). After mating, SP bound to sperm detaches from sperm, becoming free and active. Active SP then decays exponentially at rate *β*, which is connected to it’s molecular half-life. As flies with only free SP do not show a prolonged post-mating response in experiments (Peng et al., 2005), we assumed that free SP decays relatively fast so that a majority of free peptides disappear within one day (which we modelled by setting *β* = 2). The dynamics of active sex peptide are described by

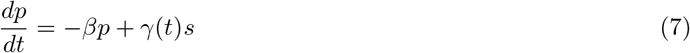

where *γ*_*t*_ is the rate at which SP is cleaved from sperm. The stock of SP attached to sperm is limited by the amount of sperm provided and it should decrease over time. We modelled this as

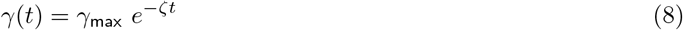

where *γ*_max_ is the initial and maximal value of *γ*, and *ζ* the rate at which it decreases towards zero. The value of *γ*_max_ is difficult to determine and is tied to the female sensitivity to SP (see (9) below). The immunofluorescence signal of the sperm bound SP disappears within five days (Peng et al., 2005) from which we assumed *ζ* = 0.6. It follows from (7) that the SP attached to sperm can be exhausted before all of sperm has been used.

Both egg production and release were constant in the model without regulation. As SP up-regulates both, they will now vary from a base level (*δ*_0_ and *σ*_0_) to a maximum value (*δ*_max_ and *σ*_max_) as SP increases. To determine the lower bound of the parameters, we considered a virgin fly and used (2) to search for values that could correspond to the observed data. Virgin flies lay small numbers (1-2 per day) of unfertilized eggs (Ashburner, 1989), and while wild and usually mated flies have between zero to approximately 50 eggs in storage (Boulétreau, 1978), the number of stored eggs might be higher among virgins. It also takes approximately three days to produce an egg (King, 1970). Because eggs are produced in the ovarioles in a conveyor like fashion, we assumed that it should take approximately three days to reach the state when each ovariole has produced one mature egg (which means around 40 eggs per female given assumption in “The model without regulation” section). We found that base egg release of *σ*_0_ = 0.03 and base egg production *δ*_0_ = 15 led to a release of approx. 2 eggs per day as virgin and reached the number of 40 eggs in storage within 3.2 days, which is within plausible range. The maximum values *δ*_max_ and *σ*_max_ were set to corresponding values of *δ* and *σ* in the model without regulation. The effects of SP on egg production and egg release are described by

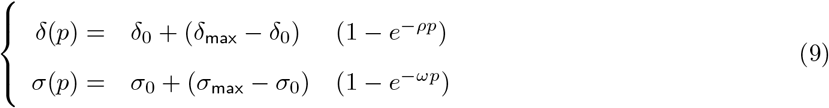

where the *ρ* and *ω* parameters can be interpreted as the sensitivity of egg production (*δ*) and release (*σ*) to SP. The higher these parameters, the less of the peptide is necessary to elicit maximal response and setting them to zero would make female unresponsive, as in the previous model. We decided that a female gains an arbitrary amount of ten free SP upon mating, and that a quantity of approximately 100 SP elicits maximal response. As we do not have any information that can distinguish the relative influence of SP on egg production or release, we treated them equally. Setting *ρ* = *ω* = 0.05 fulfils the criterion of eliciting close to maximal response with 100 SP. As the maximal number of laid eggs is observed just after mating (Klepsatel et al., 2013), we considered that this level of 100 SP should be reached early on after mating. We used *γ*_max_ = 0.5 to simulate this situation. We expect therefore an up-regulation of egg production and release to reach maximum early after mating and to decline afterwards to the base levels as SP depletes.

Unlike SP, ovulin (*v*) is provided in the form of free peptides only. We therefore modelled its dynamics as

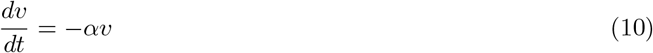

where *α* is the decay rate of ovulin. The effects of ovulin disappear after 24 hours (Herndon & Wolfner, 1995) and we propose *α* = 2, meaning that more than 85 % of ovulin is lost after first day. Ovulin is known to increase only ovulation (Herndon & Wolfner, 1995; Heifetz et al., 2000; Chapman et al., 2001), which we modelled by changing the second equation in (9) as:

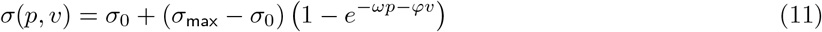

with *φ* being the egg release sensitivity to ovulin. Equation (11) implicitly assumes that in the absence of either peptide, the other can take over the stimulating role. As ovulin works through octopamine system (Rubinstein & Wolfner, 2013) and that octopamine expressing neurons may be connected to the SP sensing neurons, this is not unlikely (Häsemeyer et al., 2009; Rubinstein & Wolfner, 2013; Rezával et al., 2014; Krupp & Levine, 2014). We decided that upon mating, fly gains *v*_0_ = 100 of ovulin. As with the SP, we considered that the maximal response should be reached at this value and we thus assume that sensitivity parameter is the same as for SP (*φ* = 0.05). The larger quantity of free ovulin than that of free SP, means that we expect a strong, but short lasting effect on egg release directly after mating.

The final process in the model is the release of sperm from storage, which is increased by SP (Avila et al., 2015). We modelled this by following the same logic as that for the egg release:

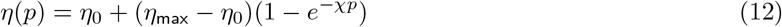

with *η*_0_ being the base level of sperm release and *η*_max_ its maximum. The maximum value is that used in the model without regulation (*η*_max_ = 3) and we set the base value to the lowest value we proposed before (*η*_0_ = 0.1). We set sensitivity of sperm release to sex peptide to *χ* = 0.05, using the same logic as for the previous sensitivity parameters - it should elicit close to maximal response initially and allow for down-regulation later on. By combining all the equations, we obtained a complete model of fly reproduction that can be regulated by the Sfps (Fig. 1, blue and black arrows).

### Regulation by seminal fluid proteins aligns the fitness components

The quantity of ovulin is maximal upon mating, while SP peaks shortly afterwards (Fig. 2C). This yields a strong up-regulation of egg production, egg release and sperm release immediately after mating (Fig. 2D) leading to an early peak in the offspring production (Fig. 2E). As sperm is gradually released, the quantity of the available SP decreases (Fig. 2C) which means that the egg and sperm release also decrease. The number of released eggs per unit of time eventually settles at the base value which corresponds to the experimental observation that females lay small number of eggs even after they stopped reproducing (Hihara, 1981; Klepsatel et al., 2013). Because the number of released eggs and sperm decline in parallel, the number of released eggs can remain longer below that of released sperm (Fig. 2F). This delays the time of sperm limitation and reduces the number of wasted eggs. Previously we suggested that the mismatch between the number of released eggs and sperm leads to the wastage of eggs and makes it impossible to simultaneously maximize the three fitness components. Here, adding the regulation by the Sfps appears to have reduced this wastage.

To test the hypothesis that regulation by the Sfps solves the trade-offs between total offspring production, time, and egg wastage we reproduced the analysis of Fig 3A-C and varied egg and sperm released rates. A technical complication arising from regulation by Sfps, is that those rates are now functions of SP and ovulin, rather than fixed parameters. Consequently, we varied the two maximal parameters of sperm and egg release (*η*_max_ and *σ*_max_, Fig. 3D-F) considering them the closest equivalent to the parameters in the model without regulation. In order to ensure the logical requirement that the base parameter values remain always below the maximal value, we changed the base values *σ*_0_ = 0.03 and *η*_0_ = 0.1, used in Fig. 2C-F, to *σ*_0_ = *η*_0_ = 0.01. The results of this analysis are shown in Fig. 3D-F.

When reproduction is regulated by SP and ovulin, the strategy of releasing sperm very slowly does not maximize the total number of offspring anymore (compare Fig. 3D to Fig. 3A). When sperm is released very slowly, peptides that stimulate the egg production and release can be exhausted before sperm, leading to the down-regulation of egg production and release, despite an abundance of sperm 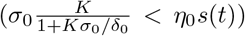. Conversely, if sperm is released too fast, some will be wasted early on, reducing the number of offspring, similar to the scenario without regulation. Taken together it means that the strategy maximizing the total number of offspring is now at an intermediate values of sperm release (3D), which value nicely matches those we considered physiologically relevant in the previous section (see Bloch Qazi & Hogdal, 2010).

Adding the Sfps has complex effects on how fast offspring are produced (Fig. 3E). Under no regulation increasing sperm release gradually accelerated reproduction (Fig. 3E). Under regulation we observe a rapid shift from slow to fast reproduction when sperm release passes a sort of threshold resulting from SP depletion.The strategy that maximizes the total offspring production lies above this “threshold” and allows for reaching 50 % of offspring in close to three days, rather than 40 days with no regulation. Regulation also reduces the number of wasted eggs (Fig. 3F), and the offspring maximizing strategy now falls in a region where the wastage of eggs is, if not minimal, in the lower range of possible values.

Overall, regulation by ovulin and SP aligns the three fitness components, and therefore almost suppresses the trade-off we identified under no regulation. Under the new optimal strategy, which corresponds to physiologically likely parameter values, offspring production is much lower than the theoretical maximum (*o* = *S*_0_). However, without regulation the evolution of females’ reproductive strategy will entirely depend on the constraints imposed by the identified trade-off. It is likely that selection in this situation will not bring females to the strategy that maximizes offspring production, but rather to one that balances the different fitness components, possibly close to the optimal strategy that we have identified with regulation.

### The depletion of sperm bound SP is key to align fitness components

Previous explanations assume that changes in the dynamics of offspring production are driven primarily by dynamics of SP. However, ovulin provides an early short term signal to release the eggs that might be partially responsible for the observed effects. To test this, we removed ovulin from the system (Fig. 4A-C). We observed no change in the pattern and the three components of fitness are aligned even without ovulin (compare Fig. 4A-C to Fig. 3D-F). Removing ovulin reduces the total number of offspring and the time to reach 50 % of offspring production (likely as a consequence of reduced fecundity). Removing ovulin also reduces the number of wasted eggs. As ovulin depletes fast, this suggests that it can increase the number of released eggs above of released sperm early in the reproductive period. Thus, although ovulin can increase total offspring production, there exists a risk of egg wastage if sperm release is too low - something that can happen just after mating when sperm is moved to storage.

**Figure 4.**
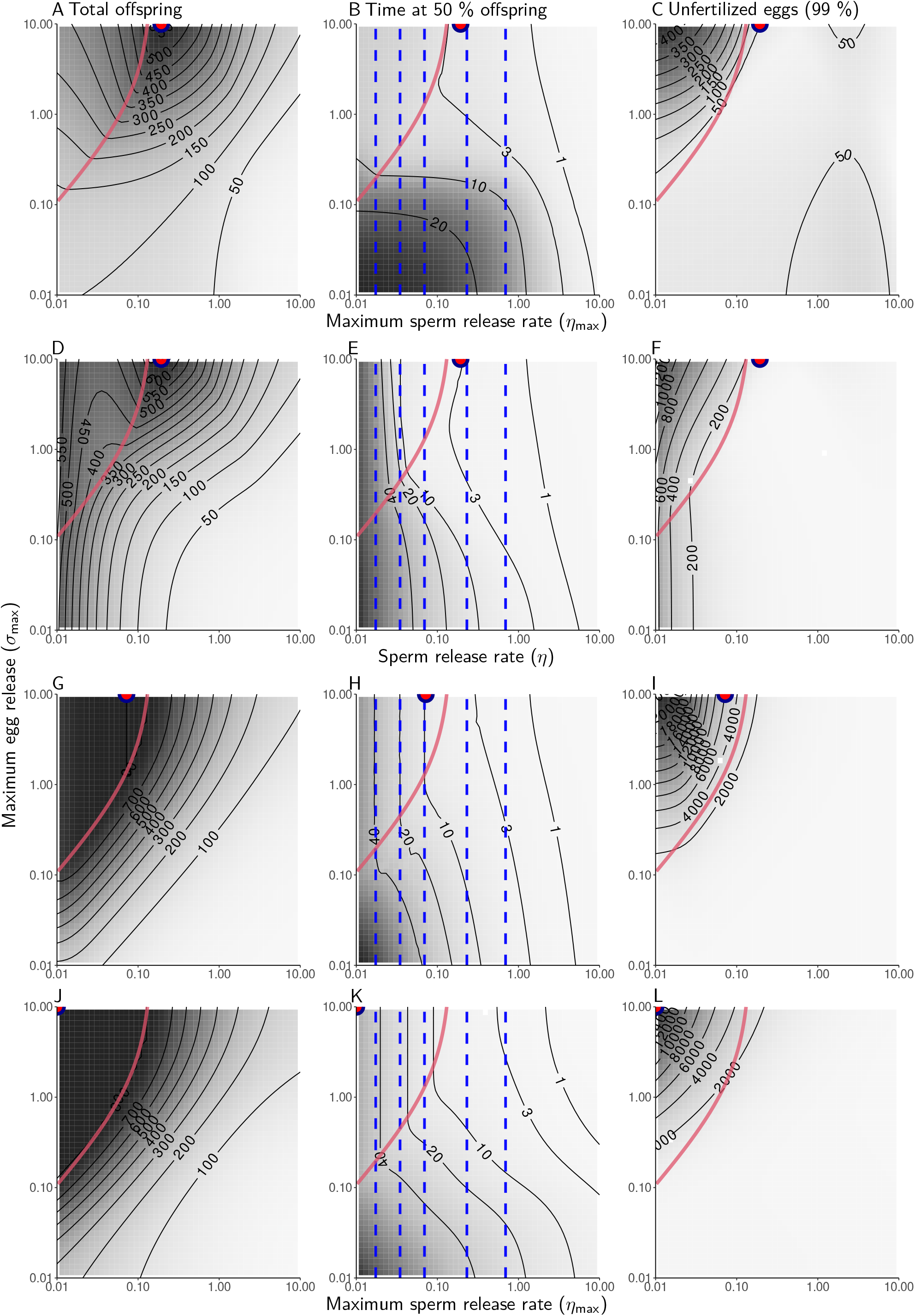
Ways regulation could lead to changes in reproduction. **(A-C) Absence of ovulin (*v* = 0)** increases speed of reproduction (B) and lowers egg wastage(C), but lowers total offspring production (A). (D-F) Sole stimulation of egg release (*δ*_0_ = *δ*_max_ = 120; *η*_0_ = *η*_max_) is sufficient to shift the optimal strategy away from slow sperm release. **(G-I) Treating sperm as signal** by preventing SP detachment and enforcing its rapid decay (*ζ* = 0; *β* = 10) is similar to cases without regulation. (J-L) A slow SP decay rate (*β* = 0.001) results in constant up-regulation of reproduction, leading to a pattern identical to no regulation (Fig. 3A-C).

We determined then that SP alone can suppress the fitness trade-offs. However, in the model SP can influence the three processes governing the reproduction. We postulated earlier that the number of released eggs is the most important element to be regulated. To test this, we restricted the effects of SP solely to the regulation of egg release and suppressed its effects on the egg production and sperm release (Fig. 4D-F). Again, this did not change the pattern and the maximal total produced offspring remains at an intermediate value of sperm release, number of wasted eggs is reduced and reproduction is similarly fast (Fig. 4E-F). We thus conclude that the effects of regulation observed in Fig. 3D-F can be explained as a consequence of the regulation of egg release by SP.

Unlike ovulin, SP produces a long term signal by its continuous release from sperm. To test if the pattern in Fig. 3D-F is a result of the prolonged signalling, we simulated a scenario where SP decay very slowly (*β* = 0.001). In this scenario SP is always present, accumulates over time and therefore does not indicate sperm availability. Under these assumptions the outcome is almost identical to the one observed without regulation (Fig. 4G-I), suggesting that slow decay of SP is similar to the constant up-regulation in the model without regulation. Thus the fact that the amount of active SP decreases over time is a key factor explaining how the three fitness components are aligned.

The active SP can decrease through molecular decay, exhaustion of sperm, or through depletion of sperm bound SP. We tested the importance of the latter mechanism by making the stock of sperm bound SP unlimited (*ζ* = 0) and letting SP decay very fast (*β* = 10). Thus the dynamics of SP exactly mirror that of sperm, as if sperm itself was used as a signal. The result is almost identical to the one without regulation (compare Fig. 4J-L to Fig. 3A-C), suggesting that egg release was continuously stimulated. This observation holds even when the sensitivity of egg release to SP (*ω*) is lowered (see Fig. S1). The depletion of sperm bound SP therefore appears to be a key mechanism explaining how regulation aligns the three fitness components. This indicates that suppressing the trade-off requires the up-regulation of egg release to decrease faster than sperm depletes.

### SP, ovulin and the potential for sexual conflict

Our results suggest that both sexes have an interest in keeping the number of sperm and eggs released closely aligned. This way the number of offspring is maximized while the number of wasted eggs is kept minimal. However, perfect synchronization is unlikely and conflict between the sexes can arise if the value of eggs differ between the sexes. Eggs are an energetic cost for the female and wastage has potential repercussions for the future reproduction. Male however gets no benefits from this future reproduction and thus is expected to prioritize efficient sperm usage. As this can be achieved by keeping number of released eggs constantly higher than released sperm release, providing a dishonest signal overestimating sperm quantity might be in male’s benefit.

We simulated divergence between signal and number of sperm by varying the number of provided sperm (*s*_0_) and the supply of SP from sperm (*γ*_max_) (Fig. 5A-C). Because we do not vary sperm and egg release, we use the base and max values of those parameters parameters as in Fig. 1B. As expected, increasing the number of sperm leads to more offspring (Fig. 5A) and a reduction in wasted eggs (Fig. 5C), and, broadly, more sperm extends reproduction, while more SP shortens it (Fig. 5B). Conversely, increasing the supply of SP leads to more offspring being produced if enough sperm is provided, but can also result in egg wastage if sperm quantity is too low. However, this has little effect on the number of offspring, and thus brings a cost to the female but no benefit to the male (Fig. 5A-C). These results suggest that SP works almost as an indicator of the number of sperm, and that there is no strong conflict between males and females over the amount of SP.

**Figure 5.**
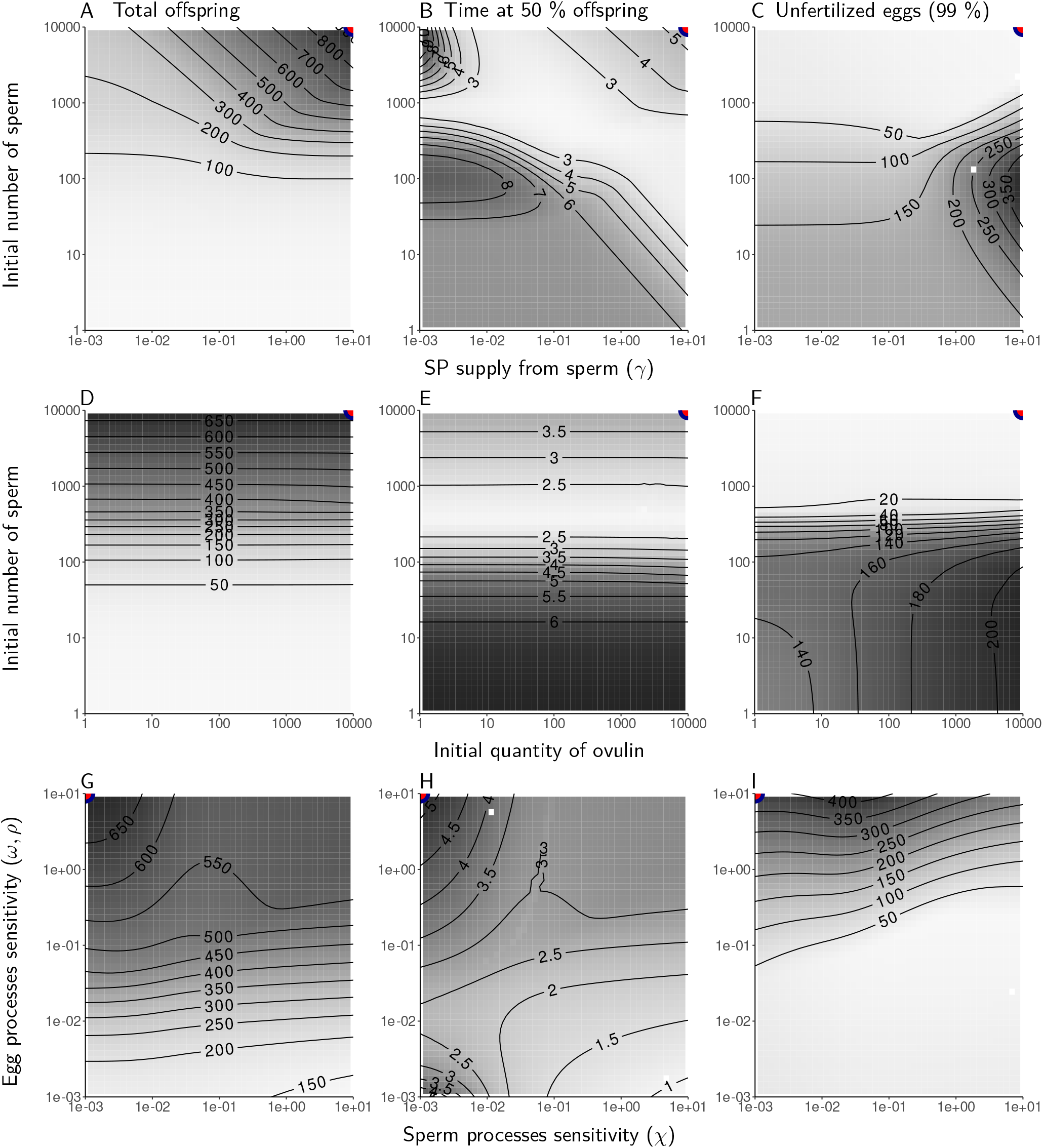
Ways males could manipulate female physiology. **(A-C) Sperm and SP mismatch.** Sperm benefit females by increasing offspring numbers (A), but it necessarily increases offspring production speed, while more SP shorten it (B). Mismatch between SP and sperm (bottom right corner, A-C) lead to egg wastage (C). **(D-F) Sperm and ovulin mismatch.** Ovulin quantity does not impact offspring production (D) and its speed (E); however, it leads to egg wastage when it exceeds sperm number (F). **(G-I) Increase SP potency.** Increased egg sensitivity to SP increases offspring production (G) but decrease its speed (H) and leads to egg wastage (I).

Aside from increasing quantity of SP, males could increase the quantity of ovulin to increase egg release. To test this, we varied the amount of ovulin and sperm that male provides (Fig. 5D-F). Here, varying ovulin quantity has almost no effect on the number of offspring (but see Fig. 5D, in contrast with Fig. 4A-C, where base sperm and egg release were low). It also does not impact the time at which 50 % of offspring is reached (Fig. 5E). However, increasing amount of ovulin leads to egg wastage if there is too few sperm provided. The conclusion it then that, similar to SP, there is probably no strong conflict between males and females over the amount of ovulin.

Another possibility to tilt female reproduction in their interest is for males to keep the amount of SP constant but to produce a highly potent variety of SP. We tested this by varying the sensitivity to SP of egg production and release (*ρ*; *ω*) against the sensitivity of sperm release (*χ*) (Fig. 5G-I). Overall, the sensitivity of sperm release has little impact on offspring, but increasing it results in faster reproduction and a slight reduction in wasted eggs (Fig. 5G-I). Increasing the sensitivity of egg production and release to SP leads to more offspring (Fig. 5G) and, because oviposition can be stimulated with lower quantity of SP, to an extended period of reproduction. This comes at the cost of an increasing amount of wasted eggs (Fig. 5G-I). The comparison of Fig. 5G and Fig. 5I suggests that the egg wastage increases faster than offspring production. This suggests in turn that a conflict might exist between males and females over the sensitivity to SP. However, when compared with the non-regulated scenario, SP seems to reduce female costs and allow for more optimal reproduction.

## Discussion

Our model provides an example of seminal fluid proteins used as signals guiding female reproduction. When egg laying is unregulated, sperm depletion results in an increasing number of unfertilized eggs being laid. If SP is used as a signal of sperm storage this wastage is reduced, while offspring production and reproductive speed is kept high. In our model, the key mechanism that allows sperm and egg release to match is depletion of SP, which diminishes the stimulating effect of SP on egg release. The up-regulation needs to wane faster than sperm is depleted to have that effect, which might explain why SP, rather than sperm, is used as signal (Eberhard, 1996; Cury et al., 2019). The binding of SP to sperm allows for the post-mating effect to last beyond short term, but also assures that stimulation ends before sperm has been depleted. SP is limited to *Drosophilinae*, but SP receptor genes have been found also in mosquitoes, moths and beetles (Yapici et al., 2008), while different components of seminal fluid can perform similar function in other insects (Eberhard, 1996; Hopkins et al., 2024). In crickets, short term increase in oviposition is driven by prostaglandins, while sperm is necessary for the long term response, potentially through similar mechanism as in *Drosophila* (Murtaugh & Denlinger, 1987; Eberhard, 1996; Larson et al., 2012). In mosquitoes, the active component, 20-hydroxyecdysone, is released over time from the mating plug and waning of the effects can be a result of the digestion of the mating plug (Gabrieli et al., 2014). Since decreasing egg production is crucial to reducing egg wastage, it is conceivable that in other species, different mechanisms evolved to ensure that egg laying does not persist beyond sperm availability.

Sfps can show a large overlap in their function (Eberhard, 1996) and in the model, both SP and ovulin can increase egg release. Redundant signals can be beneficial, especially if one acts with delay (Dore et al., 2018). Early after mating, SP binds to sperm (Misra et al., 2022) and is moved to storage, thus being potentially unavailable. We simulated this by providing less free SP than ovulin, thus most SP had to be provided by detachment from sperm. Under these conditions, ovulin increased offspring production, indicating benefits from early stimulation of egg release. Just after mating sperm release is high (Bloch Qazi & Hogdal, 2010), and it is possible that egg release due to ovulin is an adaptation to reduce sperm wastage. This assumes that female has more control over the release of eggs than sperm. However, we also observed that increased ovulin activity can increase egg wastage, something also observed experimentally (Chapman et al., 2001). Our model therefore suggests that the net effect of ovulin is determined by the balance between the increased risks of wasting eggs, and the assurance that the important, early phase of reproduction is not missed.

Sfps are often considered tools by which males can move female reproduction beyond her optimum, leading to sexual conflicts (Sirot et al., 2015). Most famously, receiving SP can reduce female lifespan in *Drosophila* (Chapman et al., 1995). However, this effect appears to be at least partially mediated through nutrition (Hopkins & Perry, 2022; Fricke et al., 2010) and might not be a general pattern for other Sfps or species, e.g. (Larson et al., 2012). As our model does not include lifespan, our conclusions about the role of Sfps in sexual conflict are limited to the implications of a dishonest, male signal on female reproductive pattern. We found that when SP potency to stimulate egg laying is enhanced, egg wastage can increase faster than offspring production. However, males obtain no benefit beyond perfect alignment of sperm and egg release. Thus the wasting of eggs would be a by product of making sure that no sperm is wasted, rather than direct exploitation. Our model predicts that the sperm-egg alignment is achieved by the continuous detachment of SP from sperm, which is largely under female control (Hopkins & Perry, 2022; Misra et al., 2022). Thus, female could easily evolve a counter adaptation to males providing too potent SP. That the degree of response is not under selection is inline with lack of coevolution between SP receptor in female and SP (Hopkins et al., 2024). Together with the lack of male benefit, ease of a female counter response, and general fitness benefits from receiving SP (Hopkins & Perry, 2022), the conflict over the effects of SP appears unlikely.

We studied females mating only once, but most flies mate multiple times (Klepsatel et al., 2013; Flatt, 2020). The results we obtained however make qualitative predictions about the implications for male-female interaction under remating. First, we observe that early after mating, egg release is often limiting, and the rate of offspring gain is at its maximum. At this stage female obtains no benefit from remating. However, after sperm becomes limiting, rate of offspring gain begins to decline and female obtains diminishing returns from the current mating. Marginal value theorem (Charnov, 1976) predicts that female should remate once that rate falls below some threshold that comes from the costs of remating. The offspring production rate can be increased by replenishing sperm and, in insects, multiple matings are often required to avoid sperm limitation (Arnqvist & Nilsson, 2000; Wang & Davis, 2006). Another way to increase that rate is the replenishment of factors stimulating sperm and egg use, which action we modelled here and that are common in insects (Eberhard, 1996; Avila et al., 2011). In crickets, the increased remating rates have been attributed to the replenishment of prostaglandins, which perform ovulin like function (Worthington et al., 2015). Males can extend the period of increased reproduction by providing the Sfps, but eventually female will benefit from remating. This leads to conflict, since, in *Drosophila*, sperm of the previous male is dumped and lost when female remates (Snook & Hosken, 2004). Seminal fluid components, including SP, have known effect to decrease female propensity to remate (Eberhard, 1996; Manning, 1967; Hopkins & Perry, 2022). Our results indicate that the remating refractory period resulting in Sfps transfer is likely to have origin in both cooperation and conflict.

## Acknowledgements

P.M. would like to thank Thomas Flatt and Sébastien Lion for helpful discussion. P.M. is supported by PhD funding through Université Toulouse 3 – Paul Sabatier and SEVAB doctoral school.

## Authors’ contributions

P.M., D.D. and J.B.F. conceptualized the study; P.M. and J.B.F. coded the model in C++; P.M. analysed the results and led the writing of the manuscript with assistance from D.D. and J.B.F.; All authors contributed critically to the drafts and gave their final approval for publication.

## Conflict of interest declaration

The authors declare they have no known financial or personal competing interests that could influence the work presented in this paper.

## Supplementary materials

Equation (1) can be solved analytically to obtain the number of eggs in storage at any time:

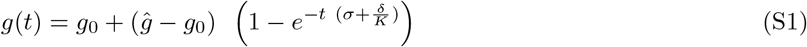

The number of stored eggs will move away from the initial number of *g*_0_, which should be zero just after the adult female fly has emerged, and will converge to the equilibrium value *ĝ*.

**Figure S1:**
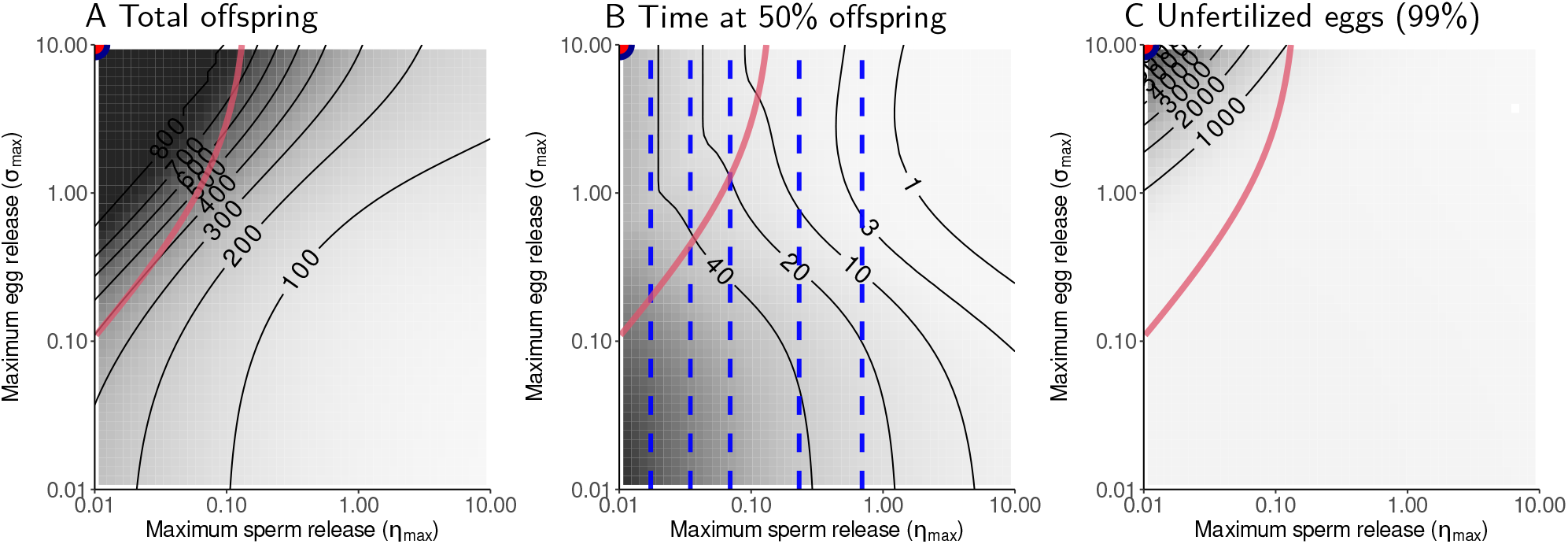
*ζ* = 0; *β* = 10; *ω* = 0.005

## Notes

### Competing Interest Statement

The authors have declared no competing interest.

